# Optic Atrophy 1 (OPA1) regulates mitochondrial resilience and injury in neonatal hypoxia–ischaemia

**DOI:** 10.64898/2026.05.21.726935

**Authors:** Claire Curel, Adam Jones, Abbe H Crawford, Ane Goikolea-Vives, Gwladys Chabrier, Gabriela Gil, Alain Oregioni, Richard Southworth, Thomas R. Eykyn, Helen B. Stolp, Irene Nobeli, Claire Thornton

## Abstract

Neonatal hypoxic–ischaemic encephalopathy is a leading cause of mortality and long-term neurodevelopmental impairment, in which mitochondrial dysfunction is central to injury progression, yet the molecular mechanisms linking mitochondrial dynamics to neuropathology remain incompletely defined.

We combined analysis of human developmental transcriptomic datasets with *in vitro* primary astrocyte models and an established neonatal mouse model of hypoxia–ischaemia (postnatal day 9, Rice–Vannucci model) to investigate the role of the mitochondrial fusion protein Optic Atrophy 1 (OPA1). OPA1 expression and processing were assessed by quantitative PCR and western blotting, mitochondrial function by imaging and Seahorse bioenergetic assays, and mitochondrial DNA content by quantitative PCR in both astrocytes and whole-brain tissue following injury.

Hypoxia–ischaemia induced rapid proteolytic processing of OPA1 in the neonatal brain and reduced OPA1 expression in astrocytes following oxygen–glucose deprivation. Genetic reduction of OPA1 in astrocytes resulted in mitochondrial fragmentation, impaired maximal respiration and spare respiratory capacity (p<0.05), and increased susceptibility to hypoxic stress (p<0.01). *In vitro*, OPA1 loss was associated with significant depletion of mitochondrial DNA (p<0.05), and mitochondrial DNA content was similarly reduced in the neonatal mouse brain 24 hours after hypoxia–ischaemia compared with controls (p<0.05). In contrast, OPA1 overexpression preserved mtDNA levels and significantly reduced *in vivo* brain tissue loss at 7 days after injury (p<0.05), while improving astrocyte survival following metabolic stress (p<0.001).

These findings identify loss of mitochondrial DNA as a previously unrecognised component of mitochondrial pathology in neonatal hypoxic–ischaemic brain injury and demonstrate that OPA1 is a key determinant of mitochondrial integrity and bioenergetic resilience. Targeting OPA1-dependent pathways may represent a novel therapeutic strategy to limit brain injury following birth asphyxia.

## Introduction

Neonatal hypoxic-ischaemic encephalopathy (HIE), a consequence of birth asphyxia, is a significant cause of mortality and morbidity worldwide, affecting 1-2 per 1000 babies in the UK and up to 26 per 1000 in low resource countries.^1^ The consequences for babies and parents are devastating; worldwide, 28% of babies with HIE will die in the neonatal period, and 24% will develop lifelong neurocognitive impairment, with significant emotional and financial burdens on the family and on society.^2^

In infants during birth and in animal models of hypoxic-ischaemic injury (HI), impaired cerebral blood flow reduces ATP production, and availability of phosphocreatine and glucose, resulting in depletion of tissue energy reserves. Reperfusion causes a transient recovery, a “latent phase”, providing a treatment window before the injury becomes irreversible. A subsequent secondary energy failure lasts from hours to days, and is the period when the majority of cell death occurs.^3^ As long-term outcome correlates with the extent of cell death (and therefore injury), treating infants during the latent phase has the potential for life-long benefit. The only currently available therapy, therapeutic hypothermia, can be implemented up to six hours after birth and therefore sits within the latent phase.^4^ Although therapeutic hypothermia can significantly improve neurological outcomes, it is not universally successful (the number needed to treat for one successful outcome is 7), and requires significant infrastructure (making it less accessible in low resource settings). Recent data from the Global Burden of Disease database (2021) reported that although mortality due to birth asphyxia is decreasing, there is an increased prevalence of neonatal encephalopathy (of which HIE is the most common cause) especially in low resource settings.^5^ These data indicate that although more babies are surviving, a lack of effective treatment options means these infants face life-long neurodevelopmental consequences and new therapies are urgently needed.^2^ Better understanding of the cellular pathology behind HI will provide novel avenues for such therapeutic development.

Mitochondrial dysfunction acts as a hub of the injury responses triggered following birth asphyxia. Mitochondrial swelling, impaired ATP production, mitochondrial reactive oxygen species and increased mitochondrial fission are characteristic of the injury^6^ and neurons are especially vulnerable due to their high energy demand.^7^ Impaired neuronal mitochondrial metabolism, architecture and calcium overload are all observed in a rodent model of neonatal HI.^8–10^ Preclinical studies using primary cells and the well-characterised rodent neonatal HI model have also identified that mitochondrial dynamics (fission and fusion) and mitophagy are impacted by HI.^11–13^ Mitochondrial fusion is mediated by the co-ordinated activity of mitofusins (MFN1/2) at the outer mitochondrial membrane and Optic Atrophy (OPA)1 at the inner membrane, whereas fission is mediated by Dynamin-related protein (DRP)1.^14^ We previously demonstrated that in primary neurons subjected to oxygen/glucose deprivation (OGD) and after HI in neonatal mice, mitochondrial fusion is impaired through increased proteolytic cleavage of OPA1, and waves of mitophagy are induced.^11,12^

Astrocytes are key regulators of cerebral metabolic homeostasis and injury responses, modulating metabolism, glutamate clearance, inflammatory signalling, and tissue repair.^15^ Astrocyte bioenergetics are impaired following HI in neonatal mice through altered glutamine and lactate metabolism and disruption of mitochondria ultrastructure. Extracellular glutamate clearance is reduced and there are increases in both astrocyte activation and production of pro- and anti-inflammatory mediators.^10,16–19^ In this study we will build on our previous neuronal work by investigating the impact of genetically down-and up-regulating OPA1 expression in primary astrocytes, evaluating bioenergetics and downstream consequences *in vitro*. In addition, we will use a mouse OPA1 transgenic model to determine whether maintaining whole-brain OPA1 expression through the critical period following neonatal HI *in vivo* alters injury progression. These data will contribute to a growing concept that protecting mitochondrial function may represent a key avenue for therapeutic intervention.

In this study, we use genetic regulation in combination with transcriptomics, *in vitro* bioenergetics and *in vivo* metabolic and injury models to investigate how OPA1 expression impacts mitochondrial function, astrocyte and rodent brain health in neonatal HI. We will investigate the downstream pathways affected by loss of OPA1 and address the hypothesis that maintaining whole-brain OPA1 expression during the critical post-injury period influences injury progression following neonatal HI *in vivo*. Our data aim to provide support for future investigations of OPA1 as a therapeutic target.

## Materials and methods

### Research ethics statement and mouse husbandry

All animal use and procedures were in accordance with the UK Animal (Scientific Procedures) Act 1986 and approved under local rules (Royal Veterinary College, King’s College London, Animal Welfare and Ethical Review Boards, London, UK). Mice were housed in individually ventilated cages at 20-24°C and at a humidity of 45-65% in a 12:12 hour dark/light cycle. Food and water were given *ad libitum*. OPA1floxed^20^ and OPA1 transgenic^21^ mouse lines were generously provided by the Scorrano Lab.

### Human gene expression analysis

Human cortical gene expression data across the lifespan was obtained from the Brainspan Atlas of the Developing Human Brain^22^ at the Allen Institute which contains data from 42 postmortem brains (524 tissue samples, 8 post-conception weeks to 40 years). Expression of genes in the motor and somatosensory cortex were retrieved. Data are expressed as Reads per Kilobase of transcript per Million mapped reads (RPKM).

### Neonatal hypoxia-ischaemia (HI)

Male and female C57BL/6J mice (WT, Charles River) and C57/Bl6 OPA1^tg^ mice (OPA1tg) were subjected to HI injury at postnatal day (P)9 using the Rice-Vannucci model of unilateral carotid artery ligation^23^ to generate ischaemia, followed by hypoxia (40-50 min) as per our previous studies (and Supplementary methods).^11,24^ For OPA1tg studies, WT littermates were used as controls. Pups were culled at various timepoints, and brains snap frozen or fixed in paraformaldehyde (PFA).

### Nucleic acid extraction and quantitative polymerase chain reaction (qPCR)

RNA from cerebral hemisphere samples (<30mg) and primary astrocytes was prepared using Direct-zol RNA miniprep kit (Zymo Research) according to manufacturer’s instructions. Quantitative RT-PCR reactions (200ng) were performed using the qPCRBIO Probe 1-Step Go Hi-ROX kit (PCR Biosystems) and Taqman primers (ThermoFisher) for expression of *Drp1* (Mm01342903_m1), *Fis1* (Mm00481580_m1), *Mff* (Mm01273401_m1), *Mfn1* (Mm00612599_m1), *Mfn2* (Mm00500120_m1), *Opa1* (Mm01349707_g1), *Mief1* (Mm00724569_m1), *Mief2* (Mm01234249_g1) and *GAPDH* (Mm99999915_g1). Gene expression changes were determined relative to GAPDH.^25^ DNA was extracted from hemisphere samples (<25mg) or cell pellets using the DNeasy Blood & Tissue Kit (Qiagen) according to manufacturer’s instructions. QPCR reactions were performed using the Primerdesign PrecisionPLUS SYBR qPCR Master Mix (Scientific Laboratory Supplies) with mouse primers for mitochondrial DNA (mMitoF: 5’-CTA GAA ACC CCG AAA CCA AA-3’, mMitoR: 5’-CCA GCT ATC ACC AAG CTC GT -3’) and nuclear DNA (B2MF: 5’-ATG GGA AGC CGA ACA TAC TG-3’ and B2MR: 5’-CAG TCT CAG TGG GGGTGA AT-3’).^26^ Changes in mtDNA were determined relative to nuclear (B2M) DNA.^27^

### Western blot

Primary astrocytes and mouse tissue protein extracts were prepared, protein lysates (50 μg) resolved by SDS-PAGE, and analysed by western blot as described previously.^28^ Proteins were detected using anti-mouse OPA1 (1:1000, Abcam ab42364), anti-mouse GAPDH (1:1000, Merck G8795) and the appropriate secondary antibody (1:10,000, LICORBio IRDye). Membranes were imaged in the LICORBio Odyssey Infrared Imaging System (LICORBio UK Ltd., Cambridge, UK) and quantified using LICOR Image Studio analysis software (v5.5, LICORBio).

### Primary astrocyte culture preparation and OPA1 knockdown

Astrocytes were prepared from male and female WT and OPA1^flx/flx^ mice at postnatal day 4-7. Brains were dissected and tissue from the same sex pooled (one pooled preparation per biological replicate). Tissue was triturated in DMEM containing 10% FBS and 1% penicillin/streptomycin. Following centrifugation (1000g, 3 min), cells were resuspended in DMEM growth medium and seeded on poly-D-lysine coated flasks (37°C, 5% CO_2_). After approximately 7 days, confluent cultures were shaken at 430rpm for 6-7 hours at 37°C to remove less adherent microglia and oligodendrocytes, and the remaining astrocytes seeded onto PDL-coated plates for experiments. For *in vitro* loss of OPA1, astrocytes from OPA1^flx/flx^ mice were treated with adenoviral GFP-Cre (AdCre) or GFP-alone (AdGFP; ViraQuest) in serum free DMEM for 2 hours at 37°C (multiplicity of infection = 200). Following removal of the viral medium, cells were incubated for 3-4 days for optimal OPA1 knockdown.

### Oxygen/Glucose deprivation (OD/OGD)

OD or OGD was performed for 3h at 37 °C using a Billups-Rothenberg hypoxia chamber flooded with 100% N_2_ as described previously.^28^ Cells were washed in PBS and returned to growth conditions prior to harvesting.

### Nuclear and Mitochondrial Staining in Live Cells

Astrocytes were incubated with nuclear (Hoechst 33342, 10 μg/mL) and/or mitochondria (Mitotracker Orange, 100 nM, ThermoFisher) stains for 30 min (37 °C, 5% CO_2_). Stained cells were washed, imaged and counted using EVOS M5000 (x4 objective) or SP8 confocal (x63 objective) (Leica Microsystems) microscopes. Mitochondrial images were preprocessed in Image J to reduce background and then analysed using the image analysis pipeline Nellie^29^ through a Napari GUI (a minimum of 10 cells per replicate).

### Mitochondrial stress test assay (Seahorse)

Cells (20,000/well) were plated on 96-well Seahorse plates (Agilent, Stockport, UK) and treated with AdCre or AdGFP virus as above. Seahorse cartridges were prepared and a Mito Stress Test (Agilent) performed according to the manufacturer’s instructions using oligomycin (1.5μM), FCCP (1.0μM) and rotenone/antimycin A (0.5μM). Following the assay, cells were stained with Hoechst (10µg/ml, 30 min), imaged and counted. Oxygen Consumption Rates (OCR) were normalized to the cell counts for each well and an average of 4-6 wells reported per experiment.

### Differential gene expression analysis

RNA from adenovirus-treated astrocytes (RNA integrity number>6, ≥ 50ng/µl, A260/A280 = 1.8-2.2) was processed by Azenta Life Sciences using PolyA selection libraries on the Illumina Novaseq platform (20 million 2×150bp paired end reads per sample). Raw fastq files were quality checked using FastQC and adapter sequence trimming and quality filtering were performed using Fastp. Counts were normalized (for PCA) and differential genes identified using the DESeq2 package in RStudio (v. 4.2.3). The Database for Annotation, Visualization and Integrated Discovery (DAVID) was used to perform gene ontology (GO) analysis and Kyoto Encyclopedia of Genes and Genomes (KEGG) pathway analysis with adjusted p-values. Figures were generated in RStudio using GGplot2.

### Histology and image analysis

Pups were terminally anaesthetised with pentobarbital at P16 (7 days post injury, i.p. 0.1ml/kg) and immediately perfused with saline via cardiac puncture. The whole brain was collected and immersion fixed in 4% paraformaldehyde for 7 days at 4°C. Samples were batch processed by a Tissue-Tek VIP® following standard protocols for dehydrating and paraffin wax embedding. Brain sections (10µm) were collected from the cerebral cortex from five independent levels 200µm apart (rostral corpus callosum through to the caudal hippocampus). Slides from each level were stained with Haematoxylin and Eosin (H&E) using standard protocols before mounting. Images were acquired on a Zeiss Axioscan slide scanner to compare infarct of the ipsilateral (HI) with the contralateral (H). Brain areas were calculated by using an ImageJ macro for semi-automation as previously described.^24^ The ipsilateral hemisphere was represented as proportion of uninjured contralateral hemisphere. Two sequential sections were analysed per level, and all images were independently analysed by two researchers, blinded to treatment and injury group. Samples with major artefacts (e.g. folding, inconsistent staining) were excluded and data were collated prior to unblinding.

### Data analysis and availability

The sample size for the *in vivo* study was between six and ten, based on a power calculation using data from previous studies^24,30,31^ which estimated a requirement of six to eight per group. This assumes a type 1 error of 0.05 with a study power of 80%. Statistical analysis was conducted using RStudio and Prism (v10, GraphPad) and unless otherwise stated, expressed as mean ± SD. Data were evaluated for normality using the Shapiro–Wilk test and then assessed using unpaired t-test, one-way/two-way ANOVA, or Kruskal Wallis test followed by *post hoc* tests, as appropriate. Test details are recorded in figure legends, p-values included in the text and p<0.05 was deemed significant. Biological replicates for primary astrocyte analyses are defined as experiments on cells derived from different litters (*n=*3-4, with a minimum of two technical replicates), and biological replicates for brain injury analysis are individual animals (mixed litters to minimise litter effects). The data and scripts that support the analyses and findings of this study are openly available in Zenodo at http://doi.org/10.5281/zenodo.19661084. Further information on Brainspan sampling and data processing can be accessed at http://www.brainspan.org. Software and version number can be found in Supplementary Table 1.

## Results

### Mitochondria dynamics genes are similarly expressed pre- and post-birth in humans

To confirm that human mitochondrial fission and fusion genes are expressed around birth at levels similar to later developmental stages, we analysed RNA expression data taken from BrainSpan^22^, an Atlas of the Developing Human Brain (Fig. 1A). Although there was gene-based variation in expression (e.g. the highest level of expression was observed for *FIS1*) there were no significant differences in the expression of *DRP1* (*P=*0.8546), *OPA1* (*P=*0.1565), *MFN1* (*P=*0.8018) or *MFN2* (*P=*0.2670) observed in the timeframe spanning term birth (21pcw-18mo, Fig. 1B).

**Figure 1:**
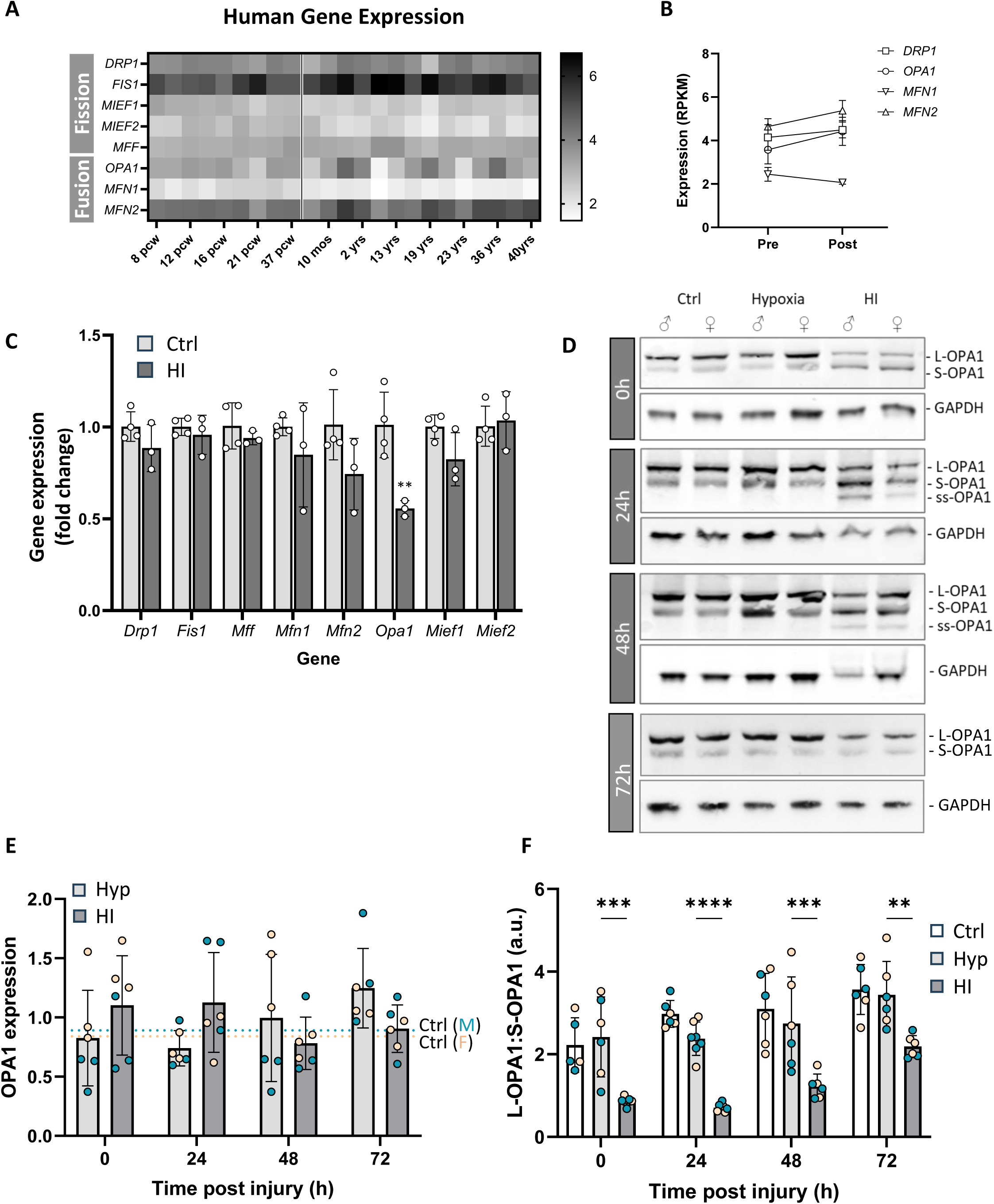
Mitochondrial fission and fusion gene expression in human brain. **A**) Heatmap showing cortical expression of selected mitochondrial fission and fusion genes across the human life span, from 8 post conception weeks (pcw) to 40 years old. **B**) Comparison of gene expression at six timepoints spanning term (around 40pcw) showing no significant differences. **C**) QPCR analysis of mRNA from Ctrl and HI brains for expression of mitochondrial dynamics genes. Data analysed by two-way ANOVA followed by Tukey *post hoc* test (*n=*3-4, \*\**P<*0.01) **D**) Protein lysates (50µg) from male and female mouse hemispheres (Ctrl, Hypoxia, HI at 0-72h post injury [50 min hypoxia]), resolved by SDS-PAGE and analysed by western blot for long (L)-OPA1, short (S)-OPA1, super short (ss)-OPA1 and GAPDH (loading control). **E**) Densitometric quantification of total OPA1 from western blots, normalised to GAPDH *n=*6 (3M,3F). **F**) Quantification of the L-OPA1:S-OPA1 ratio (including ss-OPA1 where applicable), based on densitometry. Data analysed by two-way ANOVA followed by Tukey *post hoc* test (*n=*5-7 [3-4M,2-4F] \*\**P<*0.01, \*\*\**P<*0.001, \*\*\*\**P<*0.0001).

### OPA1 is a target of neonatal hypoxic-ischaemic injury (HI) in males and females *in vivo*

We previously identified that OPA1 mRNA was depleted in primary neurons following oxygen/glucose deprivation (OGD, an *in vitro* mimic of hypoxia-ischaemia).^11^ We extended this observation by examining RNA prepared at 24h following HI in neonatal mice and found that *Opa1* was the only mitochondrial fission/fusion gene affected, decreasing expression by an average of 45% (Fig. 1C, *P=*0.0086). We next analysed OPA1 protein for changes in expression as well as sex-specific differences (Fig. 1D) as this has not been established following neonatal HI, a condition known to predominantly affect males.^32^ Total OPA1 expression was not significantly different from control for males or females although there was a trend towards increased OPA1 expression in the HI hemisphere for females compared with males at 0h (*P=*0.0509), which was not apparent at 24h (Fig. 1E). We found that long (L-) OPA1 isoforms were rapidly processed into short (S-) OPA1 forms (Fig. 1F; 0h: *P=*0.0002; 24h: *P<*0.0001; 48h: *P=*0.0003; 72h: *P=*0.0029), with a super-short (ss-) form of OPA1 visible at 24h and 48h following injury (Fig. 1D). We did not observe any sex bias in the proteolytic processing of OPA1.

### OPA1 loss sensitises astrocytes to hypoxia

We previously observed that, similar to HI in whole brain, OGD initiated OPA1 processing in neurons.^11^ As astrocyte metabolism is also impacted following neonatal HI^15^, we analysed whether astrocyte OPA1 was another target of OGD. We generated wild-type mouse primary astrocytes and validated their purity (Supplementary Fig. 1) before subjecting to OGD. We found an average 40% loss of OPA1 protein expression at 24h following OGD (Fig. 2A, *P=*0.0051) and a drop in the L-OPA1:S-OPA1 ratio (Fig. 2B, *P=*0.0073), suggesting that the well-documented negative effect of OGD on neuronal OPA1^11,13^ extends to astrocytes. To determine how OPA1 impairment affects astrocyte health, we set up a primary astrocyte model with cells generated from OPA1^flx/flx^ mice (generously provided by the Scorrano lab).^20^ Primary OPA1^flx/flx^ astrocytes were exposed to adenoviral GFP (AdGFP, control) or adenoviral GFP-tagged Cre recombinase (AdCre), which reduced OPA1 expression by over 50% (similar to OGD) without affecting viability (Supplementary Fig. 2). RNA was prepared from these cells and DESeq2 analysis used to identify differentially expressed genes (DEGs). Even with a log2 fold change cutoff of 0.3 to detect more subtle changes, we only found a total of 25 downregulated genes and 19 upregulated genes in AdCre-treated astrocytes compared with AdGFP-treated controls (adjusted p < 0.05, Fig. 2C, Supplementary Table 2). GO (Gene ontology) and KEGG (Kyoto Encyclopaedia of gene and genomes) pathway analyses were performed; pathways and GO terms with an adjusted p-value <0.05 were considered significantly enriched (Fig. 2D). There were expected significant changes in pathways associated with mitochondrial function including oxidative phosphorylation and electron transport. Of interest to this study, a response to hypoxia was also highlighted as a consequence of OPA1 knockdown. We therefore examined the effect of OGD on astrocytes with reduced OPA1 expression, compared with control. OGD induced the expected cell death in both populations of cells but there was no difference in the response to OGD between AdGFP-treated (control) and AdCre-treated (OPA1 knockdown) astrocytes (Fig. 2E) and there was no sex bias. However, when the injury was limited to hypoxia alone (oxygen deprivation, a milder injury which did not induce cell death in AdGFP astrocytes), a significant reduction in cell number was observed in astrocytes lacking OPA1 (Fig. 2F; *P=*0.0089 *vs* Ctrl AdCre; *P=*0.0098 *vs* OD AdGFP).

**Figure 2:**
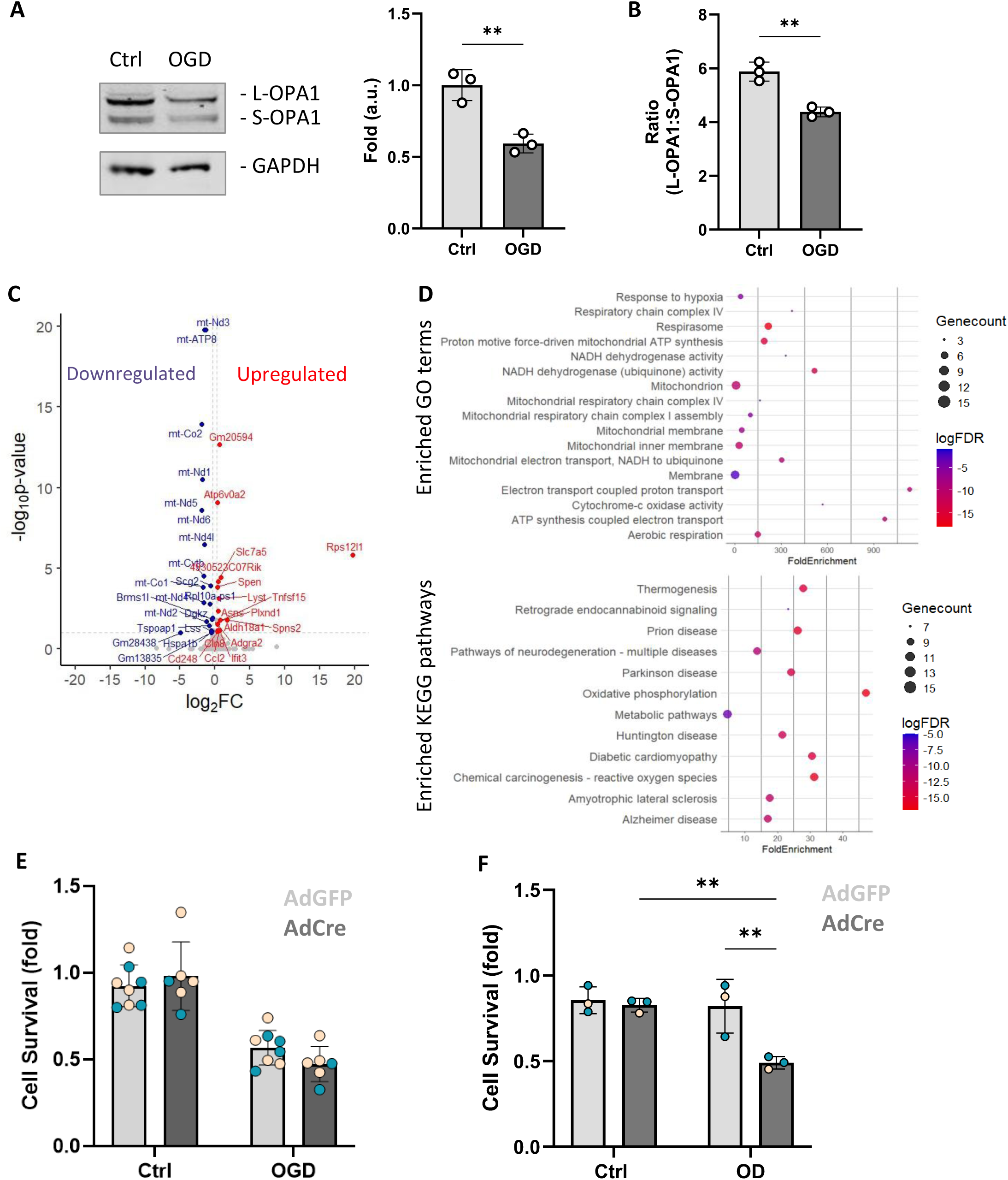
Loss of astrocyte OPA1 sensitises cells to hypoxia. **A**) Protein lysates (50µg) from primary astrocytes untreated (Ctrl) or exposed to OGD, resolved by SDS-PAGE and analysed by western blot for OPA1 expression (left panel). Densitometric analysis (right panel, normalised to GAPDH) confirmed loss of OPA1. Data analysed by unpaired t-test, \*\**P<*0.01, *n=*3. **B**) Densitometric quantification of the astrocyte L-OPA1:S-OPA1 ratio showing impact of OGD on proteolytic cleaving of L-OPA1. Data analysed by unpaired t-test \*\**P<*0.01, *n=*3. **C**) RNA was extracted from isolated primary astrocytes (Ctrl and OPA1 knockdown), sequenced on Illumina Novaseq by Azenta/Genewiz and analysed on Rstudio. Data shown as a Volcano plot of differentially expressed genes (DEGs), mapping upregulated (Red) and downregulated (Blue) genes with log_2_FC values >0.3 or <-0.3. Only DEGs with an adjusted P-value <0.05 were selected. **D**) Enriched GO terms and KEGG pathways of DEGs identified in *C*. Only pathways with a false discovery rate (FDR) < 0.05 are shown. Graphs were created using RStudio. GO: gene ontology, KEGG: Kyoto Encyclopaedia of gene and genomes. **E**) Impact of OGD on astrocyte cell survival. Adenoviral GFP-Cre (AdCre) was used to knockdown OPA1 expression with adenoviral GFP (AdGFP) used as a viral control. Data analysed by two-way ANOVA followed by Tukey *post hoc* test (*n=*6-8 [2-4M,4F], ^####^*P<*0.0001). **F**) Oxygen deprivation (OD) impairs AdCre-treated astrocytes. Data analysed by two-way ANOVA followed by Tukey *post hoc* test (*n=*3 [2M,1F], \*\**P<*0.01).

### Reduced OPA1 expression alters mitochondrial resilience

From RNA-seq analysis it was clear that mitochondrial function was significantly impacted by OPA1 loss. As expected, microscopy indicated that OPA1 knockdown induced mitochondrial fission (Fig. 3A), with a reduction in mitochondrial number (Fig. 3B, *P<*0.0001) and overall area (Fig. 3C, *P*=0.0051). This was accompanied by an increased tortuosity (increased mitochondrial curvature; Fig. 3D, *P<*0.0001), decreased branch aspect ratio (Fig. 3E, *P=*0.0052) and branch length (Fig. 3F, *P<*0.0001) indicating a reduction in the complexity of the mitochondrial reticulum. We next examined the bioenergetic consequences of OPA1 knockdown, measuring oxygen consumption rate in these cells (Fig. 3G). While mitochondrial basal respiration (*P=* 0.6092, Fig. 3H) and proton leak (*P=*0.7305, Fig. 3I) were not affected by reduction in OPA1 expression, loss of OPA1 caused a significant decrease in spare respiratory capacity (*P=*0.0126, Fig. 3J) and maximal respiration (*P=*0.0011, Fig. 3K). This indicates that reduced OPA1 expression does not affect mitochondrial respiration under basal conditions, however under cellular stress or heavy workload, OPA1 loss may impair the cell’s ability to increase energy production to meet demand due to reduced bioenergetic reserve.^33^

**Figure 3.**
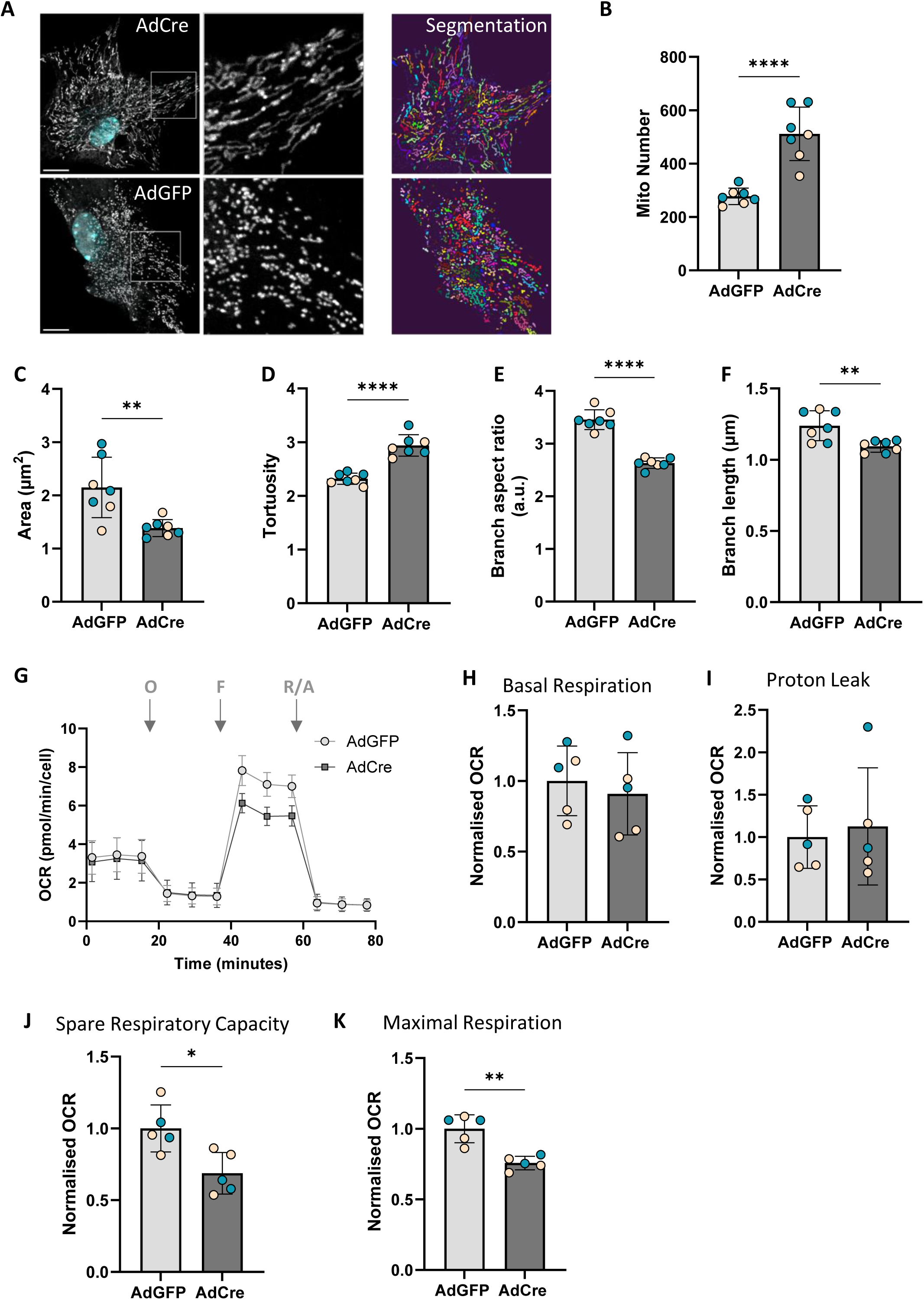
Loss of astrocyte OPA1 impairs mitochondrial function. **A**) Example image of mitotracker orange-stained mitochondria in AdGFP- or AdCre-treated astrocytes (left). Also shown is the Nellie segmentation for the same images (right). Scale bar = 10µm. **B-F**) Mitochondrial network analyses using the Nellie pipeline identified changes in mitochondrial number (**B,** \*\*\*\**P<*0.0001), mitochondrial area per cell (**C,** \*\**P<*0.01), tortuosity (**D**, \*\*\*\**P<*0.0001), branch aspect ratio (**E**, \*\*\*\**P<*0.0001) and branch length (**F,** \*\**P<*0.01). All metrics indicate mitochondrial fragmentation and reduced mitochondrial complexity following AdCre exposure. Data analysed by unpaired t-test, *n=*7 (minimum 10 cells per *n*), [4M,3F]. **G**) Representative Seahorse mito stress test of adenoviral-treated astrocytes. O: oligomycin, F: FCCP, R/A: rotenone/antimycin A. Basal respiration (**H**) and proton leak (**I**) remained unaffected but spare respiratory capacity (**J**) and maximal respiration (**K**) were impaired in astrocytes with OPA1 loss (AdCre). Data analysed by unpaired t-test, \**P<*0.05, \*\**P<*0.01 (*n=*5 [2M,3F]).

### Reducing OPA1 expression contributes to loss of mtDNA in astrocytes *in vitro* and in brain *in vivo*

In humans, mtDNA depletion is reported in conditions such as Dominant Optic Atrophy as a consequence of disease-causing OPA1 mutations.^34^ STRING analysis of RNA-seq DEGs from adenovirus-treated astrocytes (Fig. 2C, Supplementary Table 2) identified downregulation of all 13 protein-encoding genes of the mitochondrial genome (Fig. 4A). We analysed mtDNA content relative to nuclear DNA from AdCre-treated astrocytes. Interestingly, relative mtDNA content was significantly higher in male-derived astrocytes than in female astrocytes (*P=*0.0010). Despite this, mtDNA content decreased significantly in both astrocyte preparations following knockdown of OPA1 (Fig. 4B; Male: *P=*0.0485; Female: *P=*0.0321), supporting the RNA-seq data. Given that HI also results in OPA1 impairment (Fig. 1D-F), we therefore tested the hypothesis that a general reduction of mtDNA may also be a consequence. We analysed genomic DNA in mouse brain samples 24h after HI and found significantly decreased mtDNA relative to nuclear DNA content in both males and females following injury, compared with control or hypoxia alone (Fig. 4C, *P=*0.0201 *vs* H; *P=*0.0409 *vs* Ctrl).

**Figure 4.**
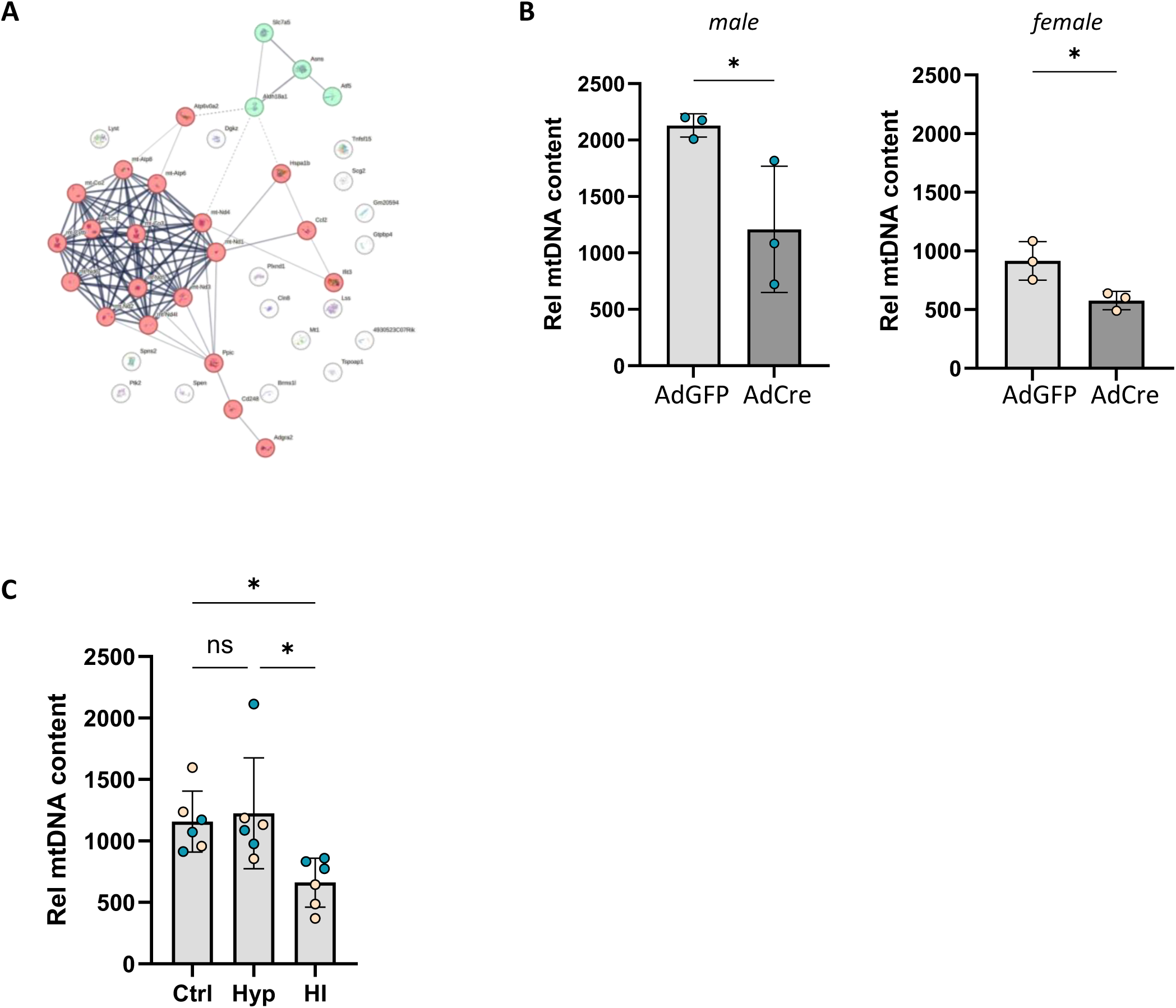
Loss of OPA1 impacts mtDNA content. **A**. STRING analysis of DEG between AdGFP and AdCre treated astrocytes with an adjusted p-value <0.1. Two clusters were identified (nodes denoted by colour) and line thickness between nodes indicates strength of supporting data. The densest clustering contains 13 protein-coding genes of the mitochondrial genome. **B**) AdCre treatment leads to a decrease in mtDNA. Expression levels of mtDNA normalised to B2M show a decrease in AdCre treated group for males (left panel) and females (right panel). Data were analysed using a unpaired t-test (*n=*3, \**P<*0.05). **C**) Genomic DNA prepared from control or HI mouse brain at 24h following injury (50 min hypoxia). MtDNA levels were significantly reduced in the HI hemisphere compared with hypoxia alone or control. Data analysed by one-way ANOVA followed by Tukey *post hoc* test, \**P<*0.05 (*n=*6 [3M,3F]).

### OPA1 overexpression provides metabolic protection against neonatal HI injury

As our data suggest that OPA1 impairment following neonatal HI impacts mitochondrial morphology, bioenergetics, and mtDNA content, we examined whether OPA1 overexpression could provide neuroprotection against injury *in vitro* and *in vivo*. We obtained the OPA1tg mouse line^21,35^, confirmed mild OPA1 overexpression (Supplementary Fig. 3) and prepared primary astrocytes from OPA1tg and WT mice. We exposed these cells to OGD, which induced an average of 40% cell death in WT astrocytes (Fig. 5A), but this was limited to 15% in astrocytes overexpressing OPA1 (Fig. 5A, *P=*0.0007). We then evaluated neuroprotection *in vivo* following neonatal HI, comparing brain tissue loss in WT and OPA1tg mice at 7 days post injury. Semi-automated blinded analyses of infarct size in H&E-stained brain sections of WT mice revealed an average of 20% loss of brain tissue in the HI hemisphere compared with hypoxia alone (Fig. 5B, *P=*0.0011). Tissue damage was significantly reduced in mice overexpressing OPA1 (Figure 5B, *P=*0.0366); there was no significant difference from uninjured mice (Fig. 5B, *P=*0.3308). Initial metabolomics data suggested that the metabolic landscape of HI-injured WT and OPA1tg mouse brain was significantly different, and there was little difference between uninjured and injured OPA1tg brain samples suggesting OPA1 overexpression may preserve metabolic stability following HI (Supplementary Methods, Supplementary Fig. 4). As loss of OPA1 *in vitro* or neonatal HI in WT mouse brain triggered loss of mtDNA (Fig. 4B,C), we investigated whether OPA1 overexpression would limit this loss. We compared levels of mtDNA at 7 days post injury (a time point matching the immunohistochemistry) and found that mtDNA in OPA1tg mice was significantly different from WT (Fig. 5C, ##*P=*0.0064). In addition, OPA1tg mice had significantly more mtDNA content than WT mice following HI injury (Fig. 5C, *P=*0.0390).

**Figure 5.**
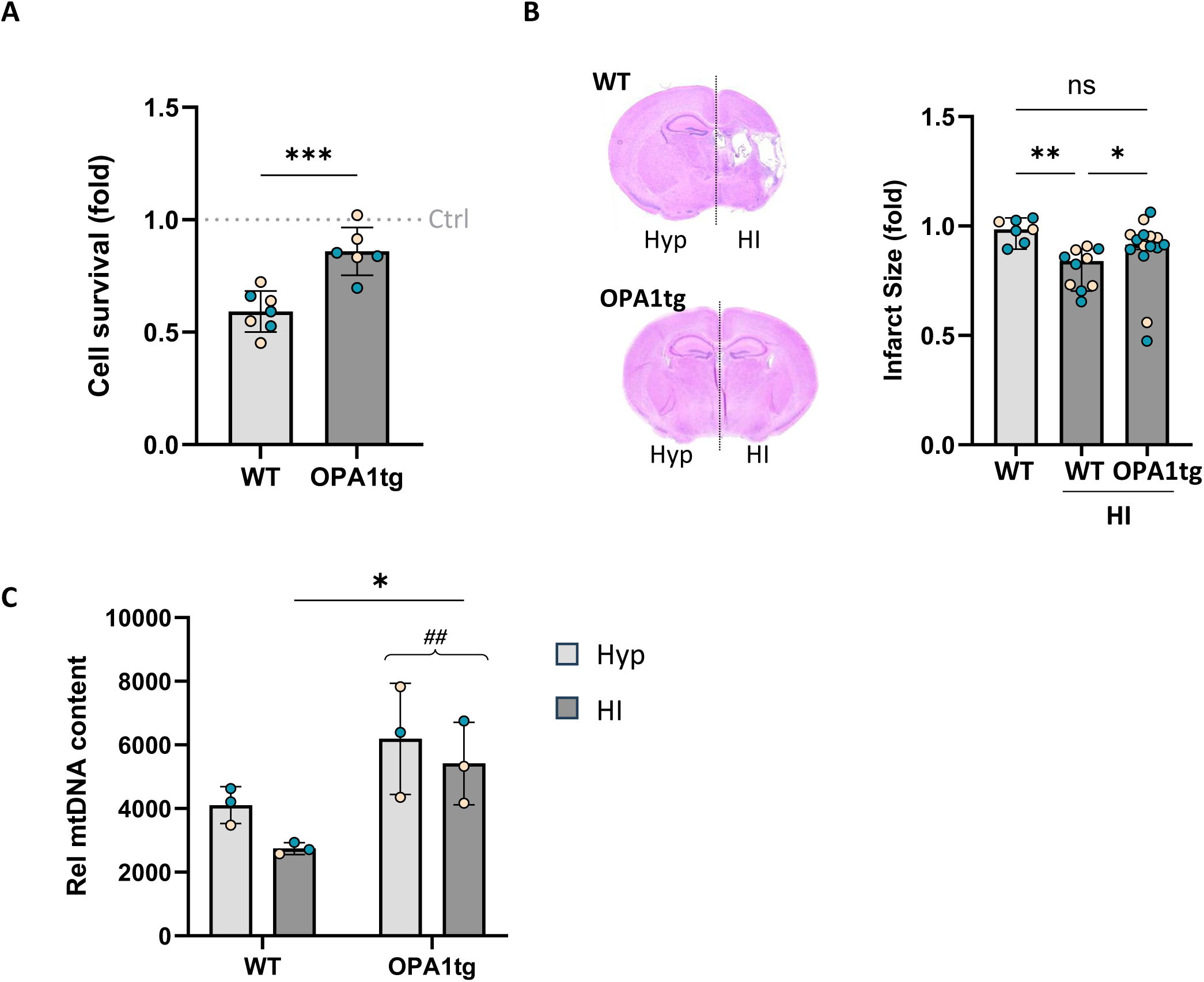
Overexpression of OPA1 is protective following HI. **A**) Primary astrocytes from OPA1tg mice were markedly protected compared with WT astrocytes, 24h after OGD. Cell number normalised to uninjured astrocytes (Ctrl) and analysed by unpaired t-test, \*\*\**P<*0.001 (*n=*6-7 [3M,3-4F]). **B**) OPA1tg mice are protected from HI. Representative brain sections from WT and OPA1tg mice (left panel) stained with H&E at 7 days following HI (40 min hypoxia). Infarct size in OPA1tg mice is indistinguishable from WT uninjured mice (ns: not significant). Quantification was performed blinded and independently by two researchers. Non-parametric data shown as median ± 95% C.I. and analysed by Kruskal-Wallis with Dunn’s *post hoc* test. \**P<*0.05, \*\**P<*0.01 (*n=*7-14 [5-8M,2-6F]). **C**) Genomic DNA prepared from WT or OPA1tg mouse brain at 7 days following injury (in line with ***B***). MtDNA content (normalised to nuclear DNA) was significantly elevated in OPA1tg mouse brain, compared with WT (##*P<*0.001) and there was a significant difference between WT and OPA1tg HI-injured mice (\**P<*0.05). Data analysed by 2-way ANOVA followed by Sidak’s *post hoc* test (*n=*3 [2M,1F]).

## Discussion

Mitochondrial dynamics play essential roles in shaping cellular resilience during development and in response to injury,^36^ yet how this machinery behaves around the time of birth and under neonatal hypoxic–ischaemic (HI) stress remains incompletely understood. Here, we identify OPA1 as a key point of vulnerability in the neonatal brain, integrating evidence from human expression datasets, *in vivo* HI, and mechanistic astrocyte models. Our findings demonstrate that OPA1 impairment contributes to hypoxia sensitivity, diminished bioenergetic reserve, and mtDNA depletion, and that increasing OPA1 expression confers significant neuroprotection both *in vitro* and *in vivo*.

Consistent expression of key genes across the perinatal period indicates that the core mitochondrial dynamics machinery remains stable during the developmental transition surrounding birth. This stability is notable given major metabolic and structural maturation in the perinatal human brain.^37,38^ Although expression varied between genes, the absence of temporal shifts supports the rationale for neonatal interventions aimed at modulating mitochondrial dynamics.

In a well-characterised rodent model of neonatal HI and compared with other mitochondria dynamics genes, OPA1 was uniquely sensitive to the injury, showing a 45% reduction in mRNA expression. Total protein levels remained comparatively stable in whole brain, although we observed decreased total OPA1 expression in astrocytes following OGD. More striking, however, was the rapid proteolytic processing of L-OPA1 into short and “super-short” forms, consistent with stress-driven activation of OMA1.^39,40^ Processed OPA1 is unable to perform efficient inner mitochondrial fusion and maintain cristae structure^41,42^ leading to mitochondrial impairment and eventual cell death. We and others previously showed aberrant OPA1 processing in neurons and in response to neonatal HI, but here we found that although 0h and 24h post-injury differences revealed minor sex-specific patterns in total OPA1 expression, the processing of OPA1 itself showed no sex bias. In humans, HI is more common in boys^32^ and this can be mirrored in rodent models, where initial absence of sex bias is reversed as the injury evolves, particularly with respect to increased inflammatory response in male mice.^43^ Although our study did not focus on sex bias, the minor nature of any sex-specific differences noted for OPA1 processing make it unlikely that OPA1 dysfunction contributes to the reported male-specific vulnerability to HI. Further studies are required to determine whether interventions based on OPA1 maintenance represent a sex-independent therapeutic target.

Pathological astrocyte activation following neonatal HI induces dysregulation of metabolic pathways and contributes to neuroinflammation.^44–46^ However, in contrast with neurons,^11–13,47^ the impact of HI on glial mitochondrial dynamics has received far less attention. Here we show that OGD induces a ∼40% reduction in astrocytic OPA1 expression and a shift toward S-OPA1, paralleling reported neuronal responses.^11,13,48^ Mimicking OGD by direct genetic OPA1 knockdown drove transcriptional changes across mitochondrial and hypoxia-responsive pathways. WT and OPA1-deficient astrocytes were similarly affected by OGD, suggesting that severe metabolic insults may obscure OPA1-dependent susceptibility. In contrast, hypoxia alone selectively killed OPA1-deficient astrocytes, revealing a previously unrecognized vulnerability under mild oxygen deprivation. These data are in line with the concept of astrocytes as the CNS sensor of PO_2_, capable of detecting physiological changes in brain oxygenation.^49^ As mitochondrial respiration is linked to astrocyte PO_2_ sensing, loss of OPA1 resulting in impaired maximal and spare respiratory capacity may sensitise these cells to hypoxic insult. Spare respiratory capacity functions as an energy reserve which mitochondria can mobilise to meet elevated energy demands caused by acute cellular stress or heavy workloads such as during HI.^50^ Astrocytes in which OPA1 is impaired may struggle to survive milder insults.

Initiated by our RNA-seq data and confirmed by qPCR, we identified that OPA1 loss results in decreased mtDNA content in astrocytes in the absence of cell death. Impaired mtDNA stability has been linked to autosomal dominant optic atrophy, a condition caused by mutations of OPA1 gene.^34,51^ OPA1 exon 4b isoform is reported to interact with the D-loop of mtDNA nucleoids, tethering them to the inner mitochondrial membrane^52,53^ and reduction of OPA1 leads to mtDNA loss. Our western blot analysis of *ex vivo* HI brain samples demonstrated that at 24h following injury, ssOPA1 is generated. As the antibodies detecting this fragment are situated at the C-terminus, and given its size, it is likely that this proteolytic product lacks the more N-terminal exon 4b, potentially releasing mtDNA. We examined this hypothesis by determining mtDNA content in HI brain samples from WT and OPA1tg mice. To our knowledge, our study is the first to show a whole-brain loss of mtDNA in WT mice at 24h following HI and that this loss is reduced in mice overexpressing OPA1. Taken together, our results suggest that the central driver of HI-mediated mtDNA depletion is OPA1 dysfunction. Loss of mtDNA is known to significantly impact cellular bioenergetics, trigger apoptosis and, in the ageing brain, contribute to the onset of neurodegeneration.^54^ In the injured developing brain, mtDNA release could act as a damage-associated molecular pattern (DAMP), contributing to the activation of pattern recognition receptors and persistent neuroinflammation.^55,56^

Finally, to combat the pathologies induced by OPA1 dysfunction, we tested OPA1 overexpression for its neuroprotective potential. We found that OPA1-overexpressing astrocytes were markedly protected from OGD-induced death. Extending this observation *in vivo*, OPA1tg mice exhibited significantly reduced infarct volumes after neonatal HI, indistinguishable from uninjured controls. These data are supported by previous observations that OPA1tg mice are protected in adult models of stroke, cardiac ischaemia/reperfusion injury and hepatocyte apoptosis.^35^ Strikingly, we also found that mtDNA levels were elevated in these mice following HI.

There are limitations and unanswered questions raised by our study. We did not determine whether loss of OPA1 in astrocytes induced an inflammatory response. Recently, pharmacological inhibition of OPA1 in neonatal HI-injured rats was linked with increased expression of components of the NLRP3 inflammasome.^57^ In a microglial cell line subjected to OGD, overexpression of OPA1 resisted upregulation of NLRP3, demonstrating a role for OPA1 in the development of neuroinflammation after HI, at least in innate immune cells.^57^ OPA1-mediated loss of mtDNA would contribute to this neuroinflammatory phenotype through its action as a DAMP.^58^ We were also not able to extend our rodent *in vivo* study to determine whether the changes in mtDNA persist long term, or whether OPA1 impairment and mtDNA loss is apparent in humans. Future work would address these issues as well as investigating the mitochondrial response in other glial cell types as currently there are limited data on the impact of neonatal HI on mitochondrial dynamics in microglia and oligodendrocytes. As OPA1 is upregulated globally in the transgenic line used here, the OPA1tg mouse model remains a significant tool in which to evaluate these glial cell types.

In summary, our findings position OPA1 as a critical determinant of mitochondrial resilience and cell survival in the neonatal brain. The vulnerability of OPA1 to HI and subsequent loss of mtDNA, combined with the substantial neuroprotective benefits of OPA1 overexpression, identifies OPA1 as a compelling therapeutic target. Given the stability of mitochondrial dynamics gene expression around birth, our data provide a mechanistic basis for developing OPA1-targeted therapies to mitigate neonatal brain injury following birth asphyxia.

## Acknowledgements

OPA1 floxed and OPA1 transgenic mouse lines were a kind gift from Prof Luca Scorrano and his lab (University of Padova, Italy), and we also appreciate their advice on the maintenance of these lines. We thank the MRC Biomedical NMR Centre at The Francis Crick Institute, which is funded by the UK Medical Research Council (CC1078), Cancer Research UK (CC1078) and the Wellcome Trust (CC1078).

## Supplementary material

Supplementary material is available at *Brain* online.

