## Supplementary material for "Optic Atrophy 1 (OPA1) regulates mitochondrial resilience and injury in neonatal hypoxia–ischaemia": Suppl.

Supplementary Data

### Mitochondrial OPA1 expression regulates the injury response to neonatal hypoxia-ischaemia

**Curel C., *et al***

#### Supplementary Methods

##### Additional Details for neonatal hypoxia-ischaemia surgery

P9 pups (min weight 4.5g, randomised in advance) were anaesthetized with 2-5% isoflurane in oxygen, with the duration of anaesthesia lasting no more than 20 minutes. The left common carotid artery of the pup was isolated and ligated with sterile surgical 0.08mm silk suture. The pups were allowed to recover for 1 hour with the dam before going through 40 (histology) – 50 (western blot) minutes of hypoxia in a warmed chamber (37°C) exposed to a humidified gas mixture composed of 10% oxygen in nitrogen. Control pups were not anaesthetized or exposed to the hypoxia chamber. Following hypoxia, all pups were returned to the dam before being culled at time points up to 7 days later via a schedule 1 method.

##### Quantitative PCR for astrocyte purity

RNA from primary cells was prepared using Direct-zol RNA miniprep kit (Zymo Research) according to manufacturer's instructions. Quantitative RT-PCR reactions (200ng) were performed using the qPCRBIO Probe 1-Step Go Hi-ROX kit (PCR Biosystems) and Taqman primers (ThermoFisher) for expression of *GFAP* (Mm01253033_m1), *Iba1*, *Tubb3*, *MBP*, *TFAM* (Mm00447485_m1) and *GAPDH* (Mm99999915_g1). Relative changes were determined for qRT-PCR relative to GAPDH (Livak and Schmittgen 2001)

##### $^1$H-NMR metabolic analysis of injured brains

Pups were culled 7 days following neonatal hypoxia ischaemia, brain tissue flash frozen and ground to a powder under liquid nitrogen. Preparation of samples was carried out as described previously (Chung *et al.* 2024). Briefly, metabolites were isolated by dual phase methanol/chloroform extraction. Aqueous extracts were resuspended in deuterated water with Trimethylsilyl propanoic acid (TSP) as an internal standard. Lipophilic extracts were resuspended in deuterated chloroform with tetramethylsilane (TMS) as internal standard. Data collection was carried out at 298K on a Bruker Avance III HD 700 MHz spectrometer equipped with a QCI cryoprobe, using randomized order. Topspin software (version 3.7) was used for data acquisition and metabolite quantification. Assignment of metabolites to their respective peaks was carried out using the ChenomX NMR Suite software. Measurement of each peak area was carried out in Topspin. Peak areas were normalised to the TSP or TMS peaks as well as the peak area sum. Heatmaps and principal component analysis (PCA) plots were generated in Matlab (vR2024b) using the 'Bioinformatics' and the 'PCA toolbox 1.5' packages. Data were autoscaled before performing PCA plots.

**Supplementary Figure 1**

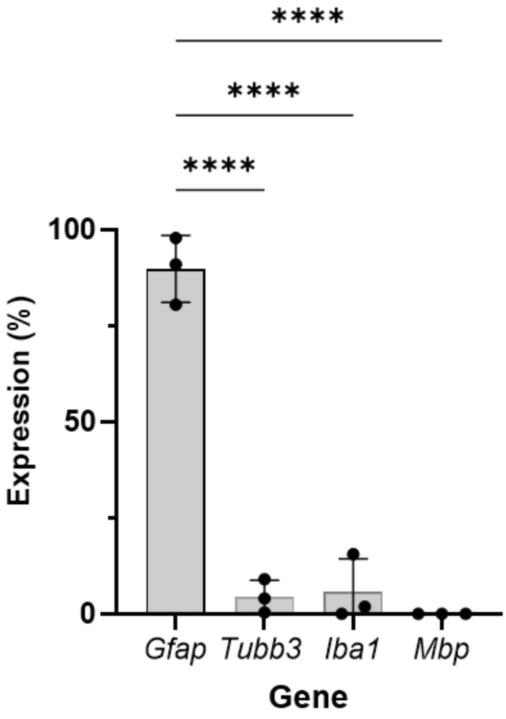


| Gene | Cell Marker | Ave Expression (%) |
|---|---|---|
| *Gfap* | Astrocytes | 89.73 |
| *Tubb3* | Neurons | 4.48 |
| *Iba1* | Microglia | 5.78 |
| *Mbp* | Oligodendrocytes | <0.01 |

**Evaluating the purity of primary astrocyte cultures.** Gene expression was analysed in primary cell culture by qRT-PCR, using markers established for astrocytes (Gfap), neurons (Tubb3), microglia (Iba1) and oligodendrocytes (Mbp). Data analysed using one-way ANOVA followed by Tukey's *post hoc* test (N=3, mean ± SD. ****P<0.0001)

**Supplementary Figure 2**

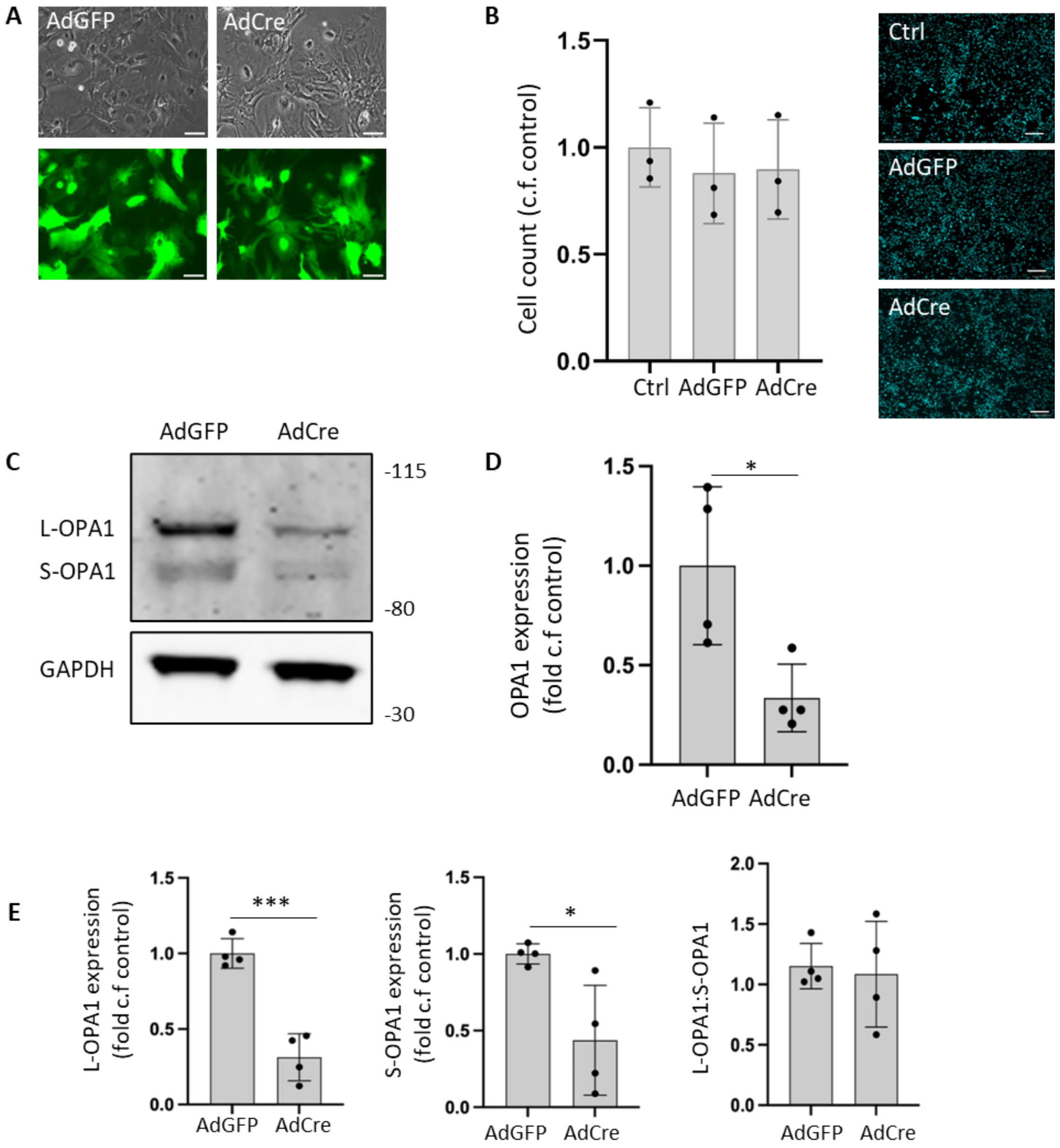


**Knockdown of OPA1 expression in primary astrocytes**. **A** Cells were treated with GFPCre or GFP for 2 hours and imaged 4 days following treatment, showing GFP expression. Scale bar = 20μm **B**. Hoechst staining and cell counting revealed no loss of viability. Scale bar = 70μm **C, D.** Protein lysates were generated from adenoviral-treated primary astrocytes taken from OPA1flx/flx mice and analysed by western blot for OPA1 expression. Quantification revealed that 4 days after exposure to Cre recombinase (200MOI), protein expression was reduced by over 50%. (*p=0.0437). GAPDH was used as a loading control. **E**. Both L-OPA1 and S-OPA1 expression were reduced (L-OPA1 ***p=0.0003, S-OPA1 *p=0.0213), but the L-OPA1:S-OPA1 ratio remained unchanged (p=0.7889) indicating that Cre-mediated knockdown altered expression of all OPA1 isoforms similarly. Data were analysed using one-way ANOVA (B) or student's t-test (D,E; N=4, mean ± SD).

**Supplementary Figure 3**

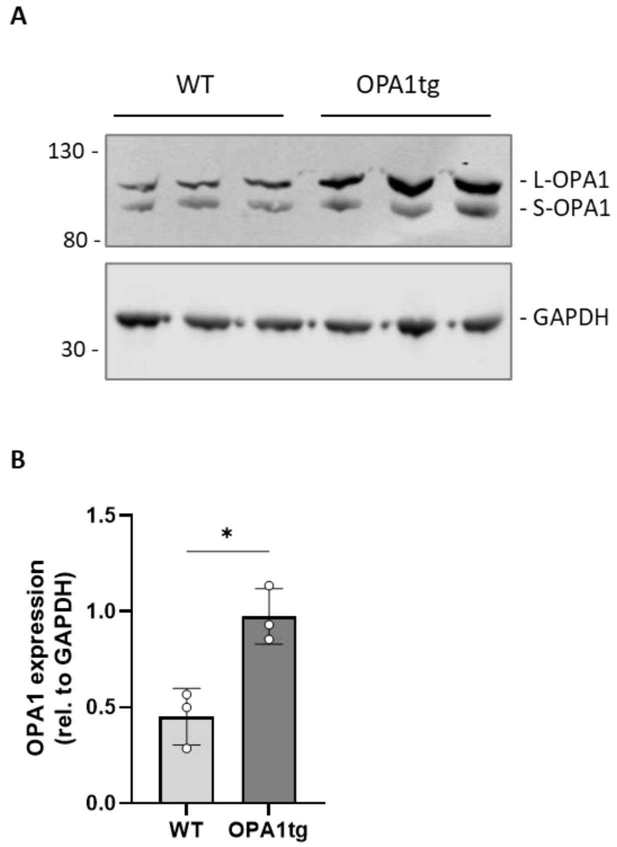


**OPA1 overexpression in astrocytes. A.** Protein lysates (50µg) were generated from primary astrocytes prepared from WT and OPA1tg mice and analysed by western blot for OPA1 expression. GAPDH was used as a loading control. **B**. Quantification revealed an approximately 2-fold overexpression in OPA1tg astrocytes compared with WT. Data were analysed by student's t-test (N=3, mean ± SD, *p=0.0198).

**Supplementary Figure 4**

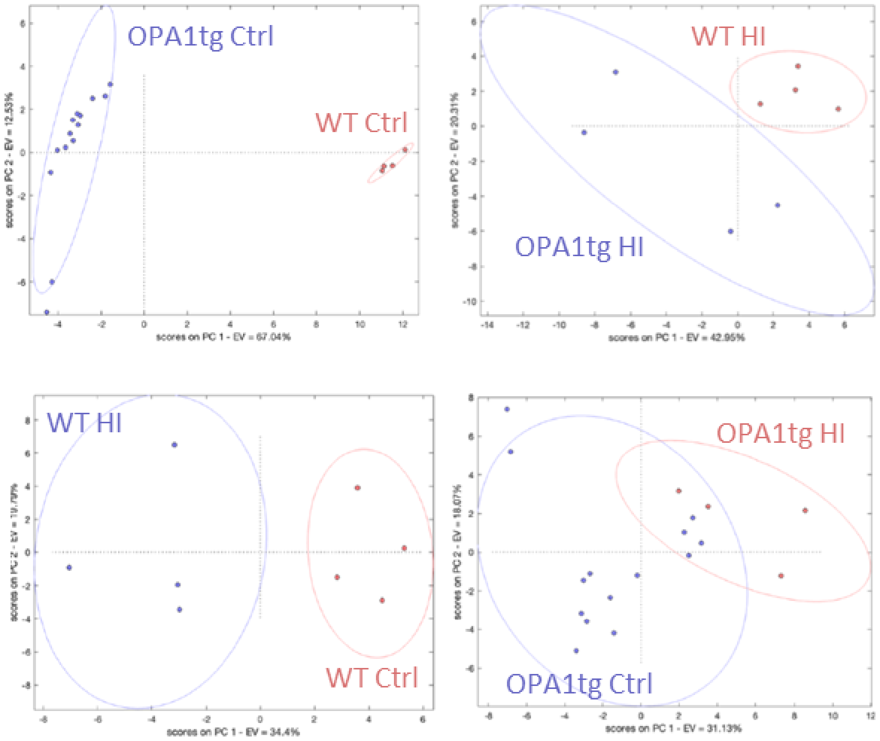


**Metabolic landscape of WT and OPA1tg mouse brain in the presence and absence of HI.** Principal component analysis (PCA) of combined lipid and metabolite data (N = 4 (WT Ctrl), 4 (WT HI), 14 (OPA1tg Ctrl), 4 (OPA1tg HI)). Data are autoscaled and ovals represent the 95% confidence regions. The variance explained by the first (PC1) and second (PC2) principal components are indicated on the axis of the scores plot. Distinct metabolic profiles are apparent for WT Ctrl vs OPA1tg Ctrl, WT HI vs OPA1tg HI and WT Ctrl vs WT HI. In contrast, the metabolic landscape of OPA1tg Ctrl vs OPA1tg HI overlap suggesting similarity.

**Supplementary Table 1: Software used for RNA-seq analysis**

| SOFTWARE | VERSION | REFERENCE |
|---|---|---|
| **FASTQC** | v0.11.9 | Andrew, S (2010) https://www.bioinformatics.babraham.ac.uk/projects/fastqc/ |
| **FASTP** | v0.23.4 | Chen *et al.* (2018) fastp: an ultra-fast all-in-one FASTQ preprocessor. *Bioinformatics,* 34**,** i884-i890. |
| **SALMON** | v1.10.0 | Patro *et al*. (2017) Salmon provides fast and bias-aware quantification of transcript expression. *Nat Methods*, 14, 417-419. |
| **RSTUDIO** | v.4.2.3 | N/A |
| **TXIMETA** | v.1.24.0 | Love *et al*. (2020) Tximeta: Reference sequence checksums for provenance identification in RNA-seq. *PLoS Comput Biol*, 16, e1007664. |
| **DESEQ2** | v.1.46.0 | Love *et al*. (2014) Moderated estimation of fold change and dispersion for RNA-seq data with DESeq2. *Genome Biol*, 15, 550. |
| **SVA** | v.3.54.0 | Leek *et al*. (2012) The sva package for removing batch effects and other unwanted variation in high-throughput experiments. Bioinformatics, 28, 882-3. |

**Supplementary table 2: Genes in which expression is altered on knockdown of Opa1**

| | Gene | | log2fold change | adj p-value |
|---|---|---|---|---|
| **Downregulated** | *mt-ATP6* | ENSMUSG00000064357 | -1.53088 | 4.00376E-24 |
| | *mt-Co3* | ENSMUSG00000064358 | -1.92079 | 4.00376E-24 |
| | *mt-ATP8* | ENSMUSG00000064356 | -1.49214 | 1.6685E-20 |
| | *mt-Nd3* | ENSMUSG00000064360 | -1.34161 | 1.6685E-20 |
| | *mt-Co2* | ENSMUSG00000064354 | -1.86833 | 1.15889E-14 |
| | *mt-Nd1* | ENSMUSG00000064341 | -1.784068 | 3.27909E-11 |
| | *mt-Nd5* | ENSMUSG00000064367 | -1.862181 | 2.49366E-09 |
| | *mt-Nd6* | ENSMUSG00000064368 | -1.883718 | 2.55398E-09 |
| | *mt-Nd4l* | ENSMUSG00000065947 | -1.53935 | 3.29214E-07 |
| | *mt-Cytb* | ENSMUSG00000064370 | -1.64636 | 2.99409E-05 |
| | *Scg2* | ENSMUSG00000050711 | -0.608389 | 0.000120805 |
| | *mt-Co1* | ENSMUSG00000064351 | -1.671936 | 0.000141231 |
| | *mt-Nd4* | ENSMUSG00000064363 | -1.569048 | 0.00130132 |
| | *Rpl10a-ps1* | ENSMUSG00000084416 | -0.711612 | 0.00168797 |
| | *Dgkz* | ENSMUSG00000040479 | -0.357436 | 0.01199424 |
| | *Brms1l* | ENSMUSG00000012076 | -0.422433 | 0.01409059 |
| | *mt-Nd2* | ENSMUSG00000064345 | -1.253975 | 0.0204015 |
| | *Tspoap1* | ENSMUSG00000034156 | -0.788453 | 0.0351328 |
| | *Mt1* | ENSMUSG00000031765 | -0.243373 | 0.0731162 |
| | *Lss* | ENSMUSG00000033105 | -0.491423 | 0.0752288 |
| | *Gtpbp4* | ENSMUSG00000021149 | -0.272751 | 0.0816031 |
| | *Hspa1b* | ENSMUSG00000090877 | -0.402506 | 0.0816031 |
| | *Gm13835* | ENSMUSG00000086922 | -0.579304 | 0.0980851 |
| | *Gm28438* | ENSMUSG00000101939 | -4.946338 | 0.0980851 |
| | *Ptk2* | ENSMUSG00000022607 | -0.246814 | 0.0996484 |

| | Gene | | log2fold change | adj p-value |
|---|---|---|---|---|
| **Upregulated** | *Gm20594* | ENSMUSG00000096887 | 0.627557 | 2.2415E-13 |
| | *Atp6v0a2* | ENSMUSG00000038023 | 0.367942 | 8.0517E-10 |
| | *Rps12l1* | ENSMUSG00000078087 | 19.69773 | 1.4726E-06 |
| | *4930523C07Rik* | ENSMUSG00000090394 | 0.86827 | 3.7564E-05 |
| | *Slc7a5* | ENSMUSG00000040010 | 0.420835 | 6.311E-05 |
| | *Spen* | ENSMUSG00000040761 | 0.371769 | 0.00014123 |
| | *Lyst* | ENSMUSG00000019726 | 0.537007 | 0.00073554 |
| | *Asns* | ENSMUSG00000029752 | 0.401522 | 0.00442715 |
| | *Atf5* | ENSMUSG00000038539 | 0.261561 | 0.0162322 |
| | *Spns2* | ENSMUSG00000040447 | 1.725948 | 0.0162322 |
| | *Tnfsf15* | ENSMUSG00000050395 | 0.791077 | 0.0162322 |

| | | | | |
|---|---|---|---|---|
| | *Plxnd1* | ENSMUSG00000030123 | 0.367241 | 0.0205132 |
| | *Aldh18a1* | ENSMUSG00000025007 | 0.383655 | 0.026503 |
| | *Adgra2* | ENSMUSG00000031486 | 0.361515 | 0.0289291 |
| | *Ifit3* | ENSMUSG00000074896 | 0.330468 | 0.0289291 |
| | *Ppic* | ENSMUSG00000024538 | 0.273138 | 0.069883 |
| | *Cln8* | ENSMUSG00000026317 | 0.385278 | 0.0732686 |
| | *Ccl2* | ENSMUSG00000035385 | 0.606176 | 0.0752288 |
| | *Cd248* | ENSMUSG00000056481 | 0.311639 | 0.0828754 |
